# MINet: a microbiome foundation model integrating microbial co-occurrence priors with abundance information

**DOI:** 10.64898/2026.11.19.767787

**Authors:** Jiaxi Zang, Hanyu Yang, Shuai Ma, Kemeng Qian, Shuyue Liu, Yong Nie, Xiao-Lei Wu

**Affiliations:** School of Mechanics and Engineering Science, Peking University, Beijing 100871, China; School of Earth and Space Science, Peking University, Beijing 100871, China; Beijing Key Laboratory of Human Microbiome Precision Medicine and Microbial Agent Development, Beijing, 100191, P. R. China; Institute of Ecology, Peking University, Beijing 100871, China

**Keywords:** autoregressive pretraining, self-supervised learning, Transformer, co-occurrence network, attention bias

## Abstract

The composition and dynamics of microbial communities are shaped by complex ecological interactions among taxa, yet most existing microbiome foundation models rely largely on the categorical identity of microbial taxa, giving limited attention to the association structure and quantitative abundance within communities. Here we present Microbial Interaction Network (MINet), a microbiome foundation model that integrates microbial co-occurrence priors with an abundance-aware mechanism. Specifically, we construct a large-scale microbial co-occurrence network from the standardized data of the Earth Microbiome Project (EMP) to provide co-occurrence-based association priors for the model, and then pretrain MINet autoregressively on more than 260,000 microbiome samples (MicroCorpus-260K). In addition to standard attention, MINet encodes z-score-normalized abundance values as input features through a two-layer multilayer perceptron, and superimposes a co-occurrence bias matrix onto the attention scores with a learnable weight, so that the attention distribution reflects both the sequence context and interactions among microorganisms. Our results show that (i) the full model consistently outperforms ablated variants with the co-occurrence bias or abundance removed in terms of pretraining loss; (ii) on the task of predicting naphthalene degradation rates for 14-strain synthetic consortia, the Transformer architecture with abundance learning significantly outperforms the vanilla Transformer and conventional machine learning methods; (iii) on downstream tasks such as biome classification and prediction of infant gut microbiota developmental stages, MINet achieves performance comparable to or better than MGM; and (iv) in generation tasks trained on data from different environments, MINet produces ecologically plausible synthetic communities whose diversity, compositional structure and co-occurrence patterns closely match those of real communities. These findings demonstrate that explicitly incorporating microbial co-occurrence priors and abundance information into pretraining can effectively enhance the representational capacity and ecological plausibility of microbiome foundation models, providing a new modelling paradigm for community-scale microbiome analysis.

## Introduction

Microbial communities inhabit nearly every environment on Earth, from host-associated body sites to oceans and soils, and contribute to host health, biogeochemical cycling, and ecosystem functioning [1–4]. The growing availability of standardized, large-scale microbiome datasets has driven the development of computational methods for characterizing and predicting community-level properties. Early approaches relied largely on conventional machine learning algorithms, including random forests, regularized regression, and support vector machines. These methods have been applied to disease-associated phenotype prediction [5–6], microbial source tracking, and environmental trait prediction. As datasets grew in size and dimensionality, supervised deep learning provided a means to learn hierarchical representations directly from microbiome profiles. Examples include EXPERT, which uses hierarchical transfer learning for cross-biome classification [7]; DeepPhylo, which incorporates phylogenetic lineage information into learned embeddings [8]; and DeepMicro, which uses autoencoders for disease-associated phenotype prediction [9]. More recently, self-supervised pretraining has been extended to microbiome analysis. MGM, for example, was pretrained on more than 260,000 unlabeled samples to learn representations that can be transferred to diverse downstream tasks [10].

Despite this progress, microbiome modeling remains challenging, particularly across heterogeneous habitats and in settings with limited labeled data. Microbiome profiles are high-dimensional, sparse, and compositional, while their associations with host or environmental phenotypes may vary across studies [11–12]. Conventional machine learning approaches often depend on task-specific feature selection or transformation and may not adequately represent higher-order relationships among taxa [13]. These challenges can increase the risk of overfitting and limit cross-study generalization when labeled datasets are small [5–6]. Although supervised deep learning reduces the need for manual feature engineering, task-specific models still require sufficient labeled data and may not transfer reliably across habitats or study populations [7–9].

Self-supervised pretraining offers a potential route to reducing this dependence on labels by learning transferable representations from large collections of unlabeled microbiome profiles [14–15]. However, the information used to construct these representations remains consequential. Models that represent taxa primarily by identity and abundance rank may lose differences in their measured abundance values that are relevant to community state. Likewise, self-attention can learn statistical associations among taxa from data, but without explicit ecological priors, it may assign weight to associations that are unstable or difficult to interpret— particularly when adaptation to a downstream task relies on few labeled samples. Incorporating abundance information and prior knowledge into pretraining may therefore improve the utility of learned representations.

Biological prior knowledge can provide an inductive bias for microbiome models by favoring representations consistent with established taxonomic, functional, or ecological structure [11–13]. One source of such knowledge is microbial co-occurrence analysis, including methods such as SparCC, SPIEC-EASI, and FlashWeave [11–12, 16]. The resulting networks summarize statistical associations among taxa; although these associations should not be equated with direct ecological interactions, they can provide a useful structural prior. For example, SIMBA-GNN uses a microbial co-occurrence network to define graph connectivity for phenotype prediction [17]. A second source is functional or phylogenetic annotation.

MicroKPNN, for instance, encodes known microbe–function relationships as sparse network connections, reducing the number of freely learned connections relative to a fully connected architecture [18]. These priors serve different purposes: functional mappings and phylogenetic lineages provide biologically defined organizational structure, whereas co-occurrence networks can be estimated from community profiles and incorporated as pairwise biases into attention. Their value for a general-purpose model, however, depends on whether associations estimated from heterogeneous environments transfer to new samples and tasks.

Here, we investigate the hypothesis that combining quantitative abundance information with a microbial co-occurrence prior during self-supervised pretraining improves the ecological relevance and downstream utility of microbiome representations. We present MINet, a microbiome foundation model with two complementary components. First, we construct a large-scale taxon co-occurrence network from standardized Earth Microbiome Project data spanning 16 environment types [2]. We incorporate this network into self-attention as a learnably weighted bias, allowing the model to use the prior without treating every inferred association as fixed or universally applicable. Second, an abundance-aware multilayer perceptron embeds z-score-normalized abundance values into taxon token representations, preserving quantitative information beyond taxon identity and rank. We pretrain MINet autoregressively on the MicroCorpus-260K dataset, which contains more than 260,000 samples, and evaluate it on synthetic consortium function prediction, hierarchical biome classification, cross-regional inflammatory bowel disease classification, infant gut microbiota developmental-stage prediction, and environment-conditioned community generation.

## Results and discussion

### Construction of the MINet model

#### Model architecture and pretraining

To enable a fair comparison with MGM, we independently rebuilt the MicroCorpus-260K corpus from MGnify records (up to June 2023), following the same collection and cleaning protocol as MGM, to serve as the pretraining corpus for MINet (Fig. 1b).This procedure retained 263,302 samples after removing low-quality or incomplete samples, yielding a vocabulary of 9,665 distinct genera. Within each sample, genera were ranked in descending order by their abundance z-scores, computed for each genus across the full corpus (see Methods), to form a "genus sequence". Because the Transformer requires inputs of a fixed length, we fixed the sequence length at 512; this specific value was chosen from the length distribution of our corpus so that 99.99% of samples fit without truncation (only the longest samples being truncated). Using these sequences, we developed MINet, a generative foundation model for microbiomes that autoregressively predicts the next genus from the preceding genera (Fig. 1c).

**Fig. 1.**
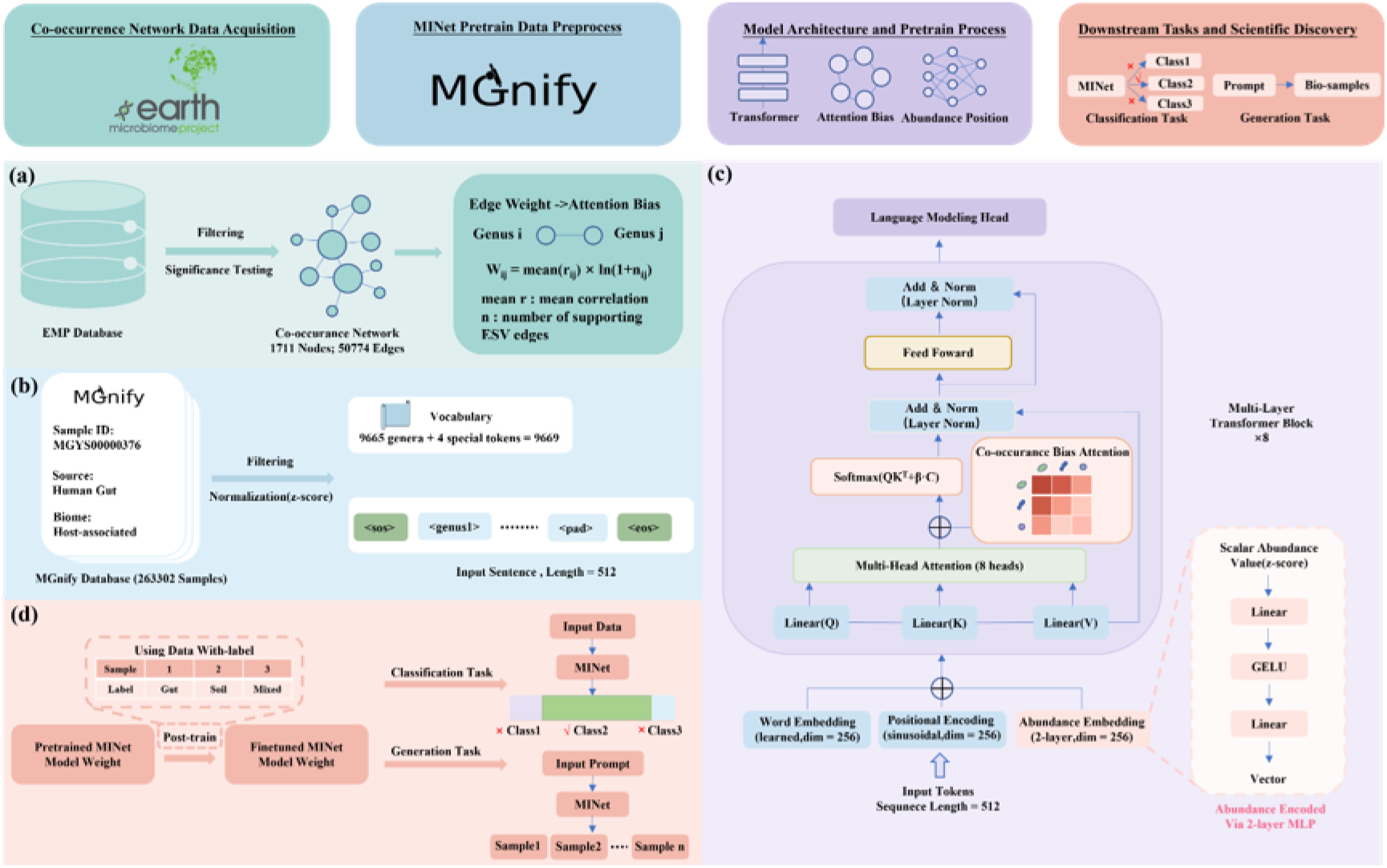
Overall workflow of MINet. (a) A large-scale microbial co-occurrence network constructed from standardized EMP data provides co-occurrence priors for the model. (b) Construction of the MicroCorpus-260K dataset: MGnify samples are cleaned and abundance-ranked to form genus sequences. (c) MINet architecture: input representations integrating abundance embeddings and co-occurrence-aware attention. (d) Transfer of pretrained representations to downstream tasks such as sample classification and community generation.

To capture both the quantitative composition and co-occurrence structure of microbial communities, MINet incorporates microbiome domain priors into its input representation and attention mechanism. Specifically, an abundance-aware multilayer perceptron (abundance MLP) encodes the continuous abundance value of each genus into an embedding, which is combined with token and position embeddings to jointly represent genus identity, ranked position and abundance. This design enables the model to distinguish differences in the dominance of the same genus across samples. This bias was derived from a global co-occurrence network that we constructed separately from standardized EMP data spanning 16 environment types (Fig. 1a). Its edges encode statistically significant pairwise associations between taxa rather than confirmed physical interactions; the full network construction is detailed in Methods. In parallel, co-occurrence-biased attention introduces a learnably weighted co-occurrence prior into the attention scores (Fig. 1c). A learnable scalar, β, adaptively balances token similarity and the co-occurrence bias, allowing the model to adjust its reliance on the prior during training.

MINet was pretrained on MicroCorpus-260K through self-supervised next-genus prediction to learn contextual representations of microbial community composition. The pretrained weights and attention scores among genera can subsequently be transferred to downstream tasks, principally sample classification and community generation across different environment types (Fig. 1d).

#### Model pretraining results and ablation experiments

To assess pretraining convergence and the individual contributions of the abundance and co-occurrence components, we trained MINet, its two ablated variants and the MGM baseline for 100 epochs under matched data splits and training settings (Methods). Their validation loss curves are shown in separate panels in Fig. 2. All four models converged stably: the validation loss fell rapidly over roughly the first 15 epochs, then entered a gradual optimization phase and continued to decline slowly through epoch 100 without rebounding.

**Fig. 2.**
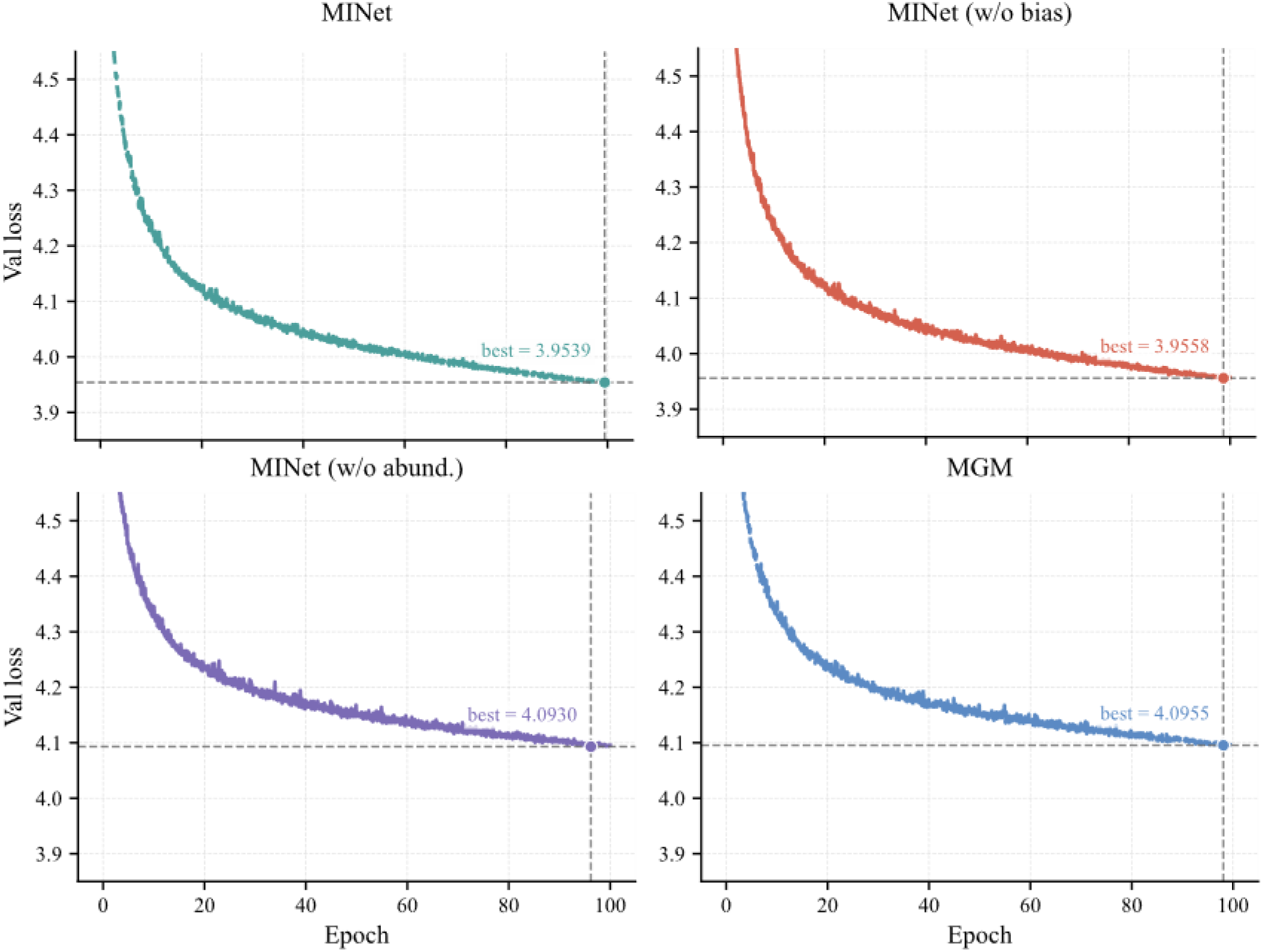
Validation loss curves of MINet, its two ablated variants and the MGM baseline over 100 epochs. Each panel corresponds to one model; the dashed line and filled circle mark the minimum validation loss reached during training (MINet, 3.9539; MINet (w/o bias), 3.9558; MINet (w/o abundance), 4.0930; MGM, 4.0955). All four models converge stably without rebound and form two tiers, with a gap of approximately 0.14 between the lower tier (MINet and MINet (w/o bias)) and the upper tier (MINet (w/o abundance) and MGM).

The validation losses separated into two clearly distinguishable tiers that persisted throughout training (Fig. 2 and Table 1). MINet reached the lowest minimum validation loss (3.9539), followed closely by MINet (w/o bias) at 3.9558; the two differed by only 0.0019. MINet (w/o abundance) and MGM formed the second tier, with minimum validation losses of 4.0930 and 4.0955, respectively, leaving a stable gap of about 0.14 between the two tiers.

**Table 1.** Minimum and final validation losses of each model.

| Model | Minimum validation loss | Final validation loss |
|---|---|---|
| MINet | 3.9539 | 3.9545 |
| MINet (w/o bias) | 3.9558 | 3.9566 |
| MINet(w/o abundance) | 4.0930 | 4.0954 |
| MGM | 4.0955 | 4.0970 |

These results point to abundance modelling as the main driver of the pretraining gain. Removing the abundance MLP raised the minimum validation loss from 3.9539 to 4.0930, bringing it back close to the MGM baseline of 4.0955. Removing the co-occurrence bias had only a marginal effect, increasing the minimum from 3.9539 to 3.9558, indicating a negligible contribution under the current setting. This limited effect may arise because the relevant information is already captured by the abundance component or the backbone, or because the pretraining objective is insensitive to what the bias encodes; its usefulness should therefore be reassessed in other tasks or at larger data scales.

### Validation and application of the MINet model

#### Prediction of degradation rates by 14-strain synthetic consortia

Before constructing MINet, we first performed a preliminary architectural validation using the synthetic-community dataset of Wang et al., in which the naphthalene degradation rates of 14 strains and their 2-, 3- and 4-member combinations were experimentally measured [19]. This experiment tested the Transformer backbone alone; the pretrained MINet model was not involved, and none of its large-scale pretraining or co-occurrence-bias components was used. Four models were compared: Transformer (rank only), in which members were ordered by their initial relative abundance and no abundance values were provided to the model; Transformer (abu.), which additionally encoded the abundance value at each position through an abundance MLP; and two tuned conventional baselines, Ridge regression and random forest (RF). The analysis asked whether the Transformer improves functional prediction over conventional methods and whether explicit abundance encoding adds predictive information beyond the ranking.

As shown in Fig. 3a, Transformer (abu.) achieved the highest Pearson correlation between predicted and measured degradation rates (r = 0.582), followed by Transformer (r = 0.512), Ridge (r = 0.469) and RF (r = 0.157). The predictions of Transformer (abu.) extended most fully along the diagonal and retained high values for highly degrading combinations. Transformer showed a similar pattern, although its predictions were less closely aligned with the diagonal. Ridge predictions were concentrated at predicted values of roughly 0.1–0.3 and systematically underestimated high-degradation combinations, whereas RF predictions showed little correspondence with the measured values.

**Fig. 3.**
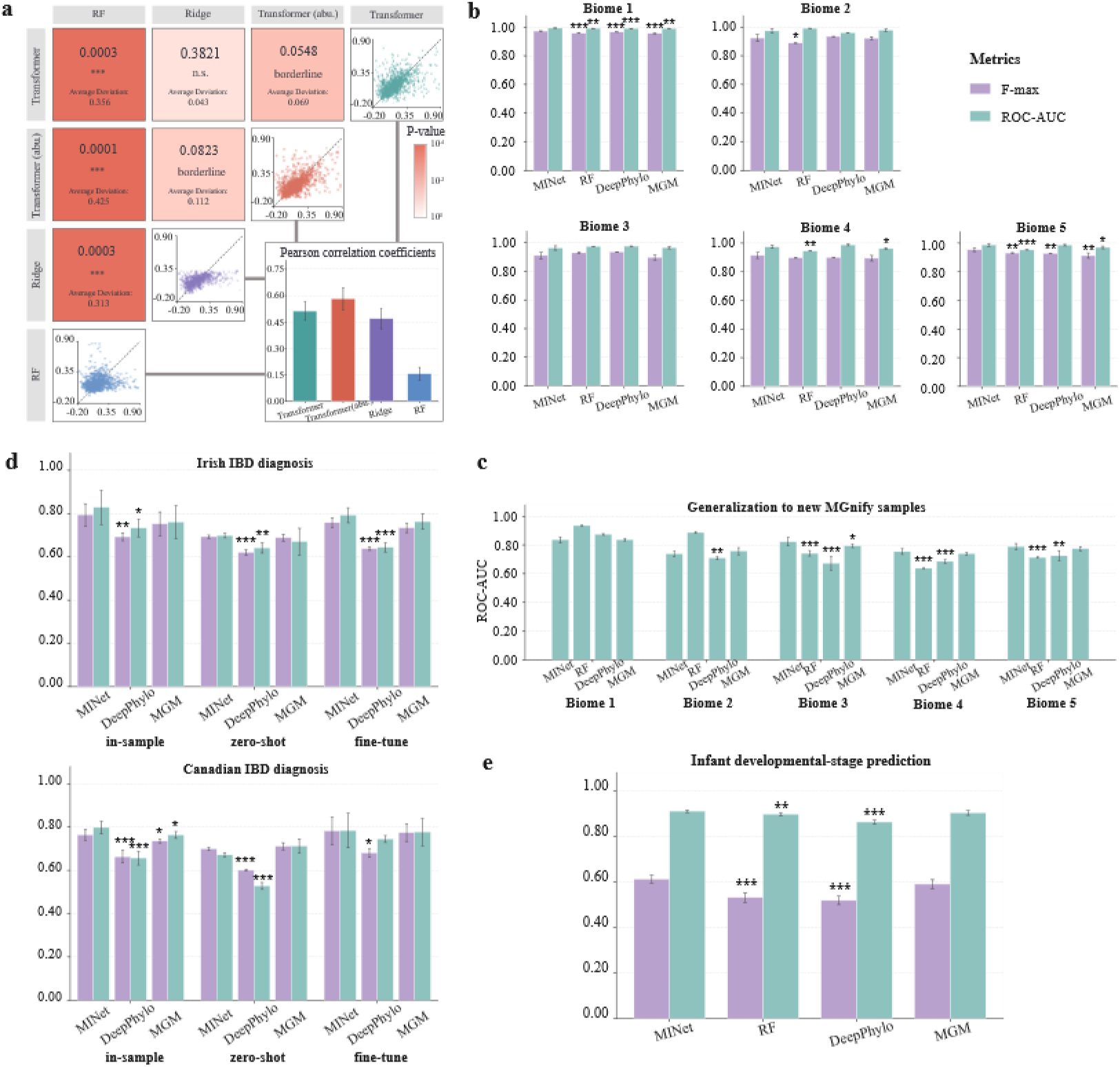
Architecture validation and downstream classification benchmarks. (a) Naphthalene degradation-rate prediction for 14-strain synthetic consortia. The upper-triangular matrix compares four models—RF, Ridge, Transformer (abu.) and Transformer. Diagonal panels show predicted versus measured degradation rates (dashed line, y = x); off-diagonal cells report two-sided paired t-tests on the per-fold Pearson correlations together with the mean difference in r; cell shading encodes the p-value, borderline denotes 0.05 ≤ P < 0.10 and n.s. denotes not significant. The enlarged lower-right panel shows mean Pearson correlations with standard deviation (SD) across five folds. (b) Ecological niche classification at five hierarchical levels (Biomes 1–5), showing F-max and ROC-AUC for MINet, RF, DeepPhylo and MGM. (c) Generalization to 75,475 MGnify samples released after the June 2023 cutoff, evaluated by ROC-AUC without parameter updates. (d) Cross-regional IBD diagnosis on Irish and Canadian cohorts under in-sample, zero-shot and fine-tune settings for MINet, DeepPhylo and MGM. (e) Infant developmental-stage classification across five time points, showing F-max and ROC-AUC for the four models. In b–e, bars are mean ± SD over five folds; asterisks denote one-sided Welch t-tests (five folds) against MINet and are shown only where MINet significantly exceeds the comparator (*P < 0.05, **P < 0.01, ***P < 0.001), and non-significant comparisons are left unlabeled.

Fig.3a summarize the cross-validation performance and pairwise comparisons. Transformer (abu.) showed a higher Pearson correlation than Transformer (mean difference +0.069, p = 0.0548) and Ridge (+0.112, p = 0.0823), although neither comparison reached p < 0.05. The difference between Transformer and Ridge was not significant (+0.043, p = 0.3821). By contrast, Transformer (abu.), Transformer and Ridge all significantly outperformed RF, with mean differences of +0.425 (p = 0.0001), +0.356 (p = 0.0003) and +0.313 (p = 0.0003), respectively. Together, these results provide preliminary support for using the Transformer architecture to predict microbial consortium function. Transformer(abu.) achieved the highest observed correlation, and the comparison with Transformer suggests that explicit abundance encoding may provide predictive information beyond abundance ranking alone. However, the comparisons with Transformer and Ridge did not reach p<0.05, so the observed improvements require cautious interpretation. This architectural validation informed the subsequent construction of the full MINet model, particularly the incorporation of an abundance MLP.

#### Performance comparison on downstream classification tasks

MINet, RF, DeepPhylo and MGM were evaluated under identical data splits, preprocessing pipelines and evaluation metrics across four classes of downstream classification tasks: ecological niche classification, generalization to newly released samples, cross-regional IBD diagnosis and infant developmental-stage prediction. Performance is reported as the mean ± standard deviation (SD) over five folds, and differences between models were assessed with one-sided Welch t-tests against MINet (Fig. 3b–e; Methods).

To assess the ability of MINet to distinguish microbial communities from different environments, we evaluated biome classification at five hierarchical levels in MGnify (Biomes 1–5), using F-max and ROC-AUC. Averaged across folds, MINet attained the highest mean F-max at Biomes 1, 4 and 5, and its mean F-max was no lower than MGM at all five levels; this statement refers to the fold-averaged means rather than to every individual fold. The strongest differences occurred at Biome 1, where MINet significantly exceeded all three comparators on both metrics (versus MGM: F-max 0.9721 ± 0.0021 vs 0.9560 ± 0.0027, P = 4.47 × 10⁻⁶; ROC-AUC 0.9934 ± 0.0008 vs 0.9904 ± 0.0012, P = 1.18 × 10⁻³), and at Biome 5 (versus MGM: F-max 0.9532 ± 0.0122 vs 0.9123 ± 0.0186, P = 2.32 × 10⁻³; ROC-AUC 0.9865 ± 0.0091 vs 0.9697 ± 0.0094, P = 1.04 × 10⁻²). At Biome 4, MINet significantly outperformed RF and MGM on ROC-AUC (versus MGM, P = 0.0342), whereas the F-max differences did not reach significance. At Biome 2, only the F-max difference from RF was significant, and no significant differences were detected at Biome 3. Overall, the gains were concentrated at the broadest level (Biome 1) and at the more specific habitat levels (Biomes 4–5), rather than being uniform across all five levels.

To evaluate generalization beyond the training data, models trained on samples released before the June 2023 cutoff were applied directly, without parameter updates, to 75,475 MGnify samples newly released between June 2023 and December 2025. At Biomes 1 and 2, no comparator was significantly lower than MINet on ROC-AUC; RF in fact attained higher mean scores (0.9354 and 0.8874, versus 0.8365 and 0.7385 for MINet). From Biome 3 onward, MINet significantly outperformed RF and DeepPhylo, and at Biome 3 it also significantly exceeded MGM (0.8227 ± 0.0295 vs 0.7938 ± 0.0125, P = 0.0478). The improvement on newly released samples was therefore concentrated at Biomes 3–5 and was not consistent across all five levels.

To assess classification across geographical regions, we evaluated two independent IBD cohorts from Ireland and Canada under three settings: in-sample, five-fold evaluation within the target cohort; zero-shot, direct application of a model trained without any target-region data; and fine-tune, further adaptation of the zero-shot model on a small target-region subset. MINet significantly outperformed DeepPhylo in nearly every setting and cohort on both F-max and ROC-AUC. Against MGM, the only significant differences occurred in the Canadian in-sample setting (F-max 0.7634 ± 0.0239 vs 0.7353 ± 0.0082, P = 0.0280; ROC-AUC 0.7981 ± 0.0287 vs 0.7634 ± 0.0157, P = 0.0270); all other MINet–MGM comparisons were not significant.

For every model, in both cohorts and on both metrics, zero-shot scores were lower than in-sample scores, and the MINet–MGM differences were not significant, with MGM scoring slightly higher on individual metrics, including ROC-AUC in the Canadian cohort. Fine-tuning largely restored performance toward the in-sample level. These results indicate that the advantage of MINet depended on the evaluation setting: significant gains were observed in some within-region comparisons and consistently against DeepPhylo, whereas a significant advantage in zero-shot cross-regional transfer was not established.

Using the longitudinal cohort of Roswall et al. and following earlier infant gut microbiota research, we evaluated developmental-stage classification across five time points: birth, 4 months, 12 months, 3 years and 5 years [20–21]. This multiclass task has imbalanced class distributions and was evaluated with F-max and ROC-AUC. MINet achieved the highest scores on both metrics (F-max 0.6117 ± 0.0173; ROC-AUC 0.9102 ± 0.0056), and its advantage over RF and DeepPhylo was significant, whereas the differences from MGM did not reach significance (F-max 0.5902 ± 0.0194; ROC-AUC 0.9031 ± 0.0103).

Across the four task classes, the performance of MINet varied with the task and evaluation setting. It significantly outperformed the comparators at selected biome levels—most clearly at Biomes 1 and 5—and, on newly released samples, at Biomes 3–5 against RF and DeepPhylo. In IBD diagnosis, MINet was consistently superior to DeepPhylo and significantly better than MGM in the Canadian in-sample setting, while in infant developmental-stage classification it significantly exceeded RF and DeepPhylo. The differences from MGM were not significant in several settings. Together, these results support the downstream classification performance of MINet, while identifying zero-shot cross-regional transfer without target-region adaptation— where no significant advantage over MGM was established—as a specific remaining limitation.

#### Comparison of community generation performance

To evaluate environment-conditioned microbial community generation, we retrieved 26,334 MGnify samples by querying the database against nine environment labels: agriculture, forest, grassland, groundwater, hot spring, lake, ocean, sediment and wetland. The retrieved samples contained 2,304 genera, all within the 9,665-genus pretraining vocabulary. Starting from the full pretrained checkpoint, we fine-tuned a conditional generator in which an environment-label token was prepended to each sequence, so that generation was explicitly conditioned on the target habitat. Because the generator emits ordered genus sequences without quantitative abundances, we trained a separate abundance reconstructor following the ranking– reconstruction framework of MGM [10]. This reconstructor was an independent model with its own parameters, trained exclusively on real training samples and never on generated sequences, and it was not updated jointly with the generator; this separation ensures that abundance assignment cannot compensate for errors made by the generator. Environment-conditioned sequences were produced with nucleus sampling (top-p = 0.9), mapped to abundance vectors by the reconstructor, and renormalized so that each synthetic community summed to 1. For the primary comparison, MINet and MGM each generated 500 communities per environment under identical decoding settings, giving 4,500 generated communities per model.

Generated communities were assessed with 13 metrics organized into five groups: diversity and sparsity (Shannon diversity, Simpson diversity and entropy-based sparsity), network topology (density, average degree and modularity of the co-occurrence network), distribution similarity (cosine similarity, Spearman correlation and mean absolute error (MAE) of abundance vectors), classification and ecological validity (environment-classification ROC-AUC and biomarker coverage), and sequence length (mean and median length). Each metric was computed per environment against the real samples of that environment and then aggregated across the nine environments. Because MINet and MGM were evaluated on the same nine environments, their per-environment values were treated as paired observations and compared using two-sided Wilcoxon signed-rank tests, which use the exact distribution at n = 9. Fig. 4 presents the metric-level comparison, and exact per-environment values are provided in Supplementary Table S1.

**Fig. 4.**
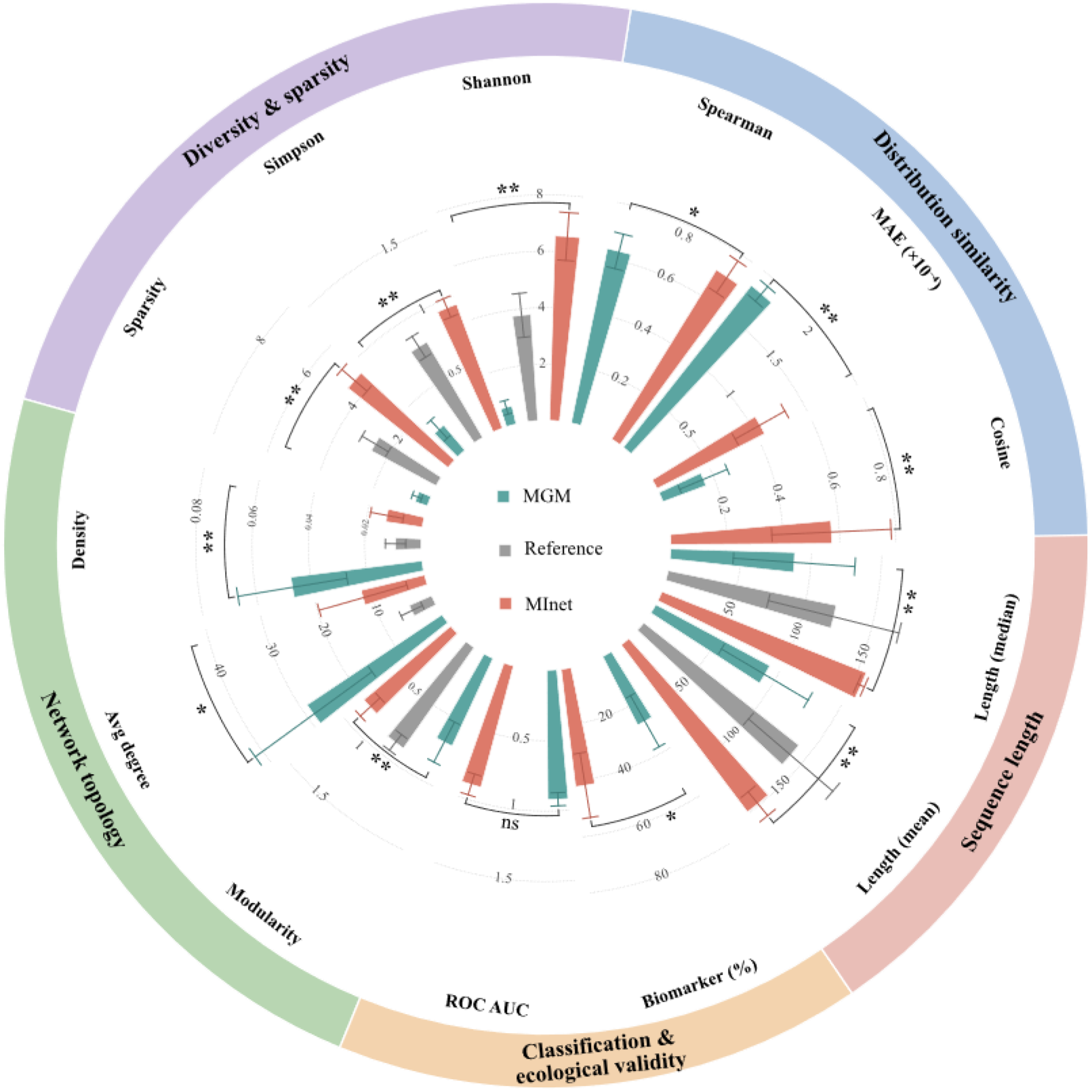
Comprehensive comparison of MINet and MGM across 13 generation-evaluation metrics. Radial bars show the mean value across the nine environments (error bars, SD) for MGM (teal), the real-data reference (grey) and MINet (coral); each radial axis uses its own scale, labelled around the outer ring, and metrics are grouped into five categories on the outer annulus. Brackets and asterisks indicate two-sided Wilcoxon signed-rank tests between MINet and MGM across the nine paired environments (*P < 0.05, **P < 0.01); n.s., not significant. Twelve of the 13 metrics differ significantly, with ROC-AUC the only exception.

MINet and MGM diverged in opposite directions from the real reference. Real communities had a mean Shannon index of 3.77 ± 0.78 across environments; MINet overestimated it (6.55 ±0.85), whereas MGM underestimated it (0.66 ±0.25). The same pattern held for Simpson diversity (MINet 0.94 ± 0.07; MGM 0.23 ± 0.09; real 0.79 ± 0.09) and for entropy-based sparsity (MINet 4.54 ±0.59; MGM 0.46 ±0.17; real 2.62 ±0.54), with all three differences significant (P < 0.01, Fig. 4). Although MINet overshot the reference on every measure, its values remained far closer to the real range than MGM’s, whose communities were consistently dominated by a small number of genera.

The two models also differed sharply in network topology. MGM markedly overestimated connectivity, producing networks denser and more highly connected than the real networks (density 0.047 ± 0.020 vs. 0.009 ± 0.004 for the real reference; average degree 28.83 ± 12.83 vs. 4.33 ±2.11). MINet’s networks were closer to the reference on both measures (density 0.013 ± 0.006; average degree 11.61 ± 8.10), and the differences between the two models were significant (P < 0.05). For modularity, MINet was close to the reference (0.81 vs. 0.87) and MGM fell below it (0.69), again a significant difference (P < 0.01). The large standard deviations for average degree indicate that these effects were heterogeneous across environments rather than uniform.

MINet reproduced the real abundance distributions more closely. Its cosine similarity (0.57 ±0.21 vs. 0.16 ±0.09) and Spearman correlation (0.71 ±0.07 vs. 0.63 ±0.06) were significantly higher, and its MAE (1.07 × 10⁻⁴ vs. 1.83 × 10⁻⁴) was significantly lower, a 41% reduction in absolute abundance error (all P < 0.05; Fig. 4).

Ecological validity presented a mixed picture. The difference in environment-classification ROC-AUC was not significant (0.892 ± 0.077 for MINet vs. 0.918 ± 0.047 for MGM; n.s., Fig. 4), and MGM in fact scored higher in several environments, including hot springs (0.9795 vs. 0.9550) and the ocean (0.9805 vs. 0.9657); MINet was higher in others, such as agriculture and forest. By contrast, MINet retained significantly more environment-associated taxa: mean biomarker coverage—the overlap of the top 50 biomarkers between generated and real communities—was 42.0 ± 11.6% for MINet versus 27.3 ± 10.4% for MGM (P < 0.05).

Finally, the models diverged in sequence length. MGM generated shorter communities than the real reference (mean length 92.0 ± 35.7 and median 88.2 ± 43.3, vs. real means of 140.3 and 122.0), whereas MINet’s lengths were close to and slightly above the reference (mean 148.7 ± 10.9; median 156.2 ± 2.7); both differences were significant (P < 0.01).

To test whether these results depended on the number of generated samples, we repeated the evaluation at six generation levels: 500, 1,000, 2,000, 5,000, 10,000 and "same", where "same" equalled the number of real training samples in the environment. The six distribution-level metrics—Shannon and Simpson diversity, sparsity, cosine similarity, Spearman correlation and MAE—were essentially unchanged across the five fixed levels, with mean coefficients of variation below 2%. The three network metrics were more sensitive to generation size and approached the real reference most closely at the "same" level. The complete metric-by-level curves are provided in Supplementary Fig. S1.

To visualize how network topology moves as generation size changes, we represented density, modularity and average degree simultaneously in ternary plots, one per environment (Fig. 5). Because the three metrics differ in scale and each environment spans a different value range, the axes were scaled independently within each panel, and each generation configuration was mapped to a point whose position reflects all three metrics; arrows connect the points in order of increasing generation size. In environments with few training samples (hot spring, 662 samples; grassland, 1,367; groundwater, 1,505), increasing the generation size moved the points progressively away from the real network (star), whereas the "same" point stayed very close to it. The opposite trend occurred in two large-sample environments (sediment, 4,984; ocean, 7,359), where increasing the generation size moved the points progressively toward the real network, and the smallest fixed size (500) was the most distant. Agriculture and forest did not follow either pattern: their points fluctuated without a clear monotonic direction, which may reflect the strong intrinsic heterogeneity of soil samples. The remaining environments (wetland and lake) showed no clear monotonic trend, but their "same" points were likewise among the closest to the real network. Across all nine environments, "same" was the configuration that most consistently landed near the real network.

**Fig. 5.**
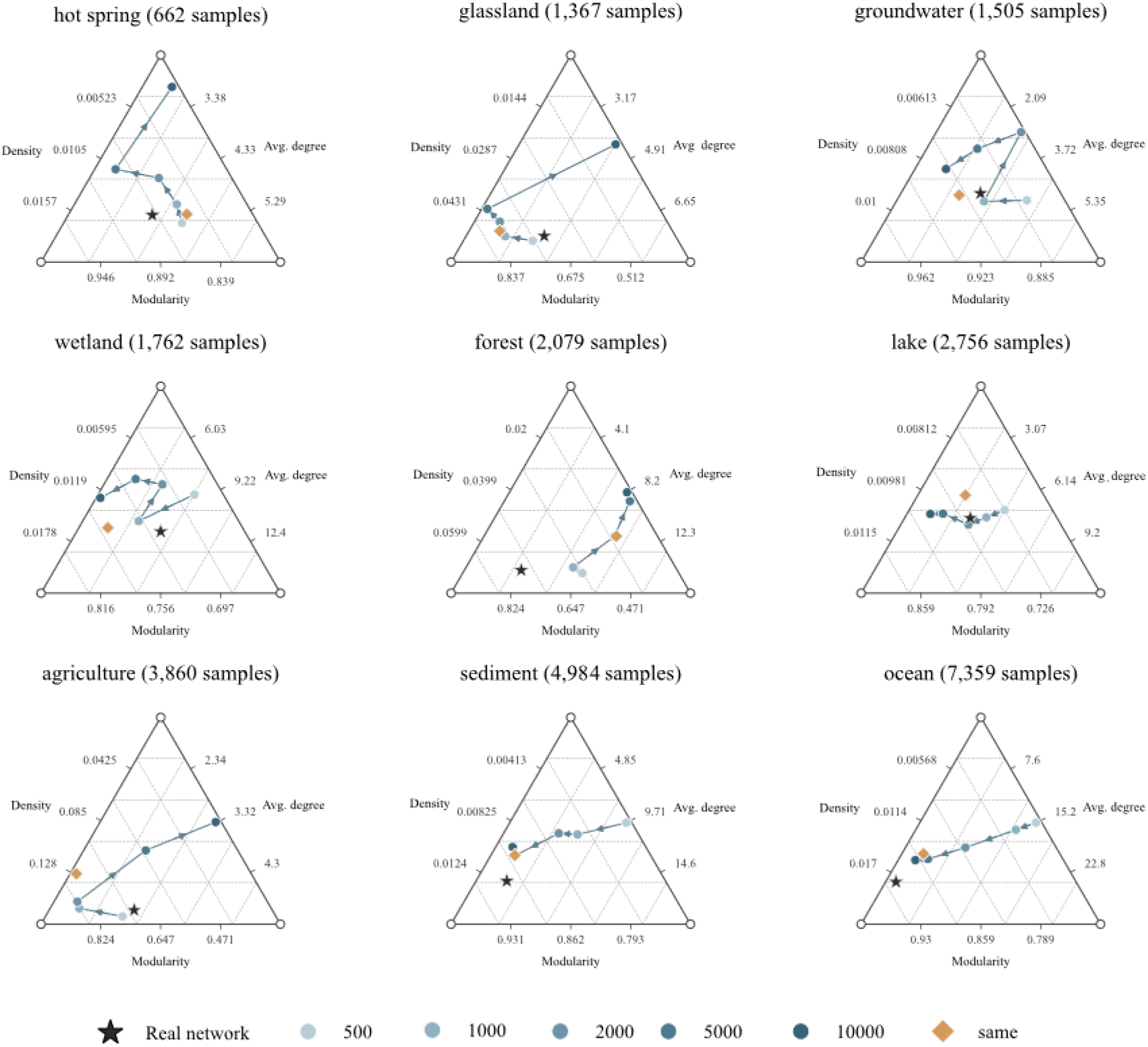
Ternary representation of co-occurrence network topology across generation sizes. Each panel corresponds to one environment (real sample count in parentheses). The three axes encode network density (top), modularity (bottom left) and average degree (bottom right), scaled independently per environment. The black star marks the real network; circles mark generated communities at five fixed sizes (500–10,000, light to dark); the orange diamond marks the "same" configuration, whose sample count equals the real training count. Arrows indicate the direction of increasing generation size. Points move away from the real network with increasing generation size in small-sample environments and toward it in large-sample environments, while the "same" point remains the configuration most consistently close to the real network.

We quantified these departures using the absolute relative deviation, |generated − real|/real, averaged over the six generation levels for each metric and environment (Fig. 6, heatmap), and further averaged across the three metrics to obtain a mean relative deviation per environment (Fig. 6, bars). Mean relative deviation was smallest in groundwater (0.19), hot spring (0.37) and wetland (0.40), and largest in ocean (2.33), forest (1.50) and agriculture (1.03), broadly increasing with the number of training samples even though the relationship was not strictly monotonic—for example, lake (2,756 samples) deviated less than forest (2,079). The non-monotonic behaviour of forest is consistent with its ternary trajectory in Fig. 5: forest soils are highly heterogeneous environments, and this strong environmental heterogeneity can introduce fluctuations in the inferred network topology rather than a smooth sample-size response. The heatmap shows that average degree contributed the largest deviations, most strikingly for the ocean (575%), with density next (253% in agriculture and 242% in forest), whereas modularity was reproduced relatively well in all environments (all cells below 42%).

**Fig. 6.**
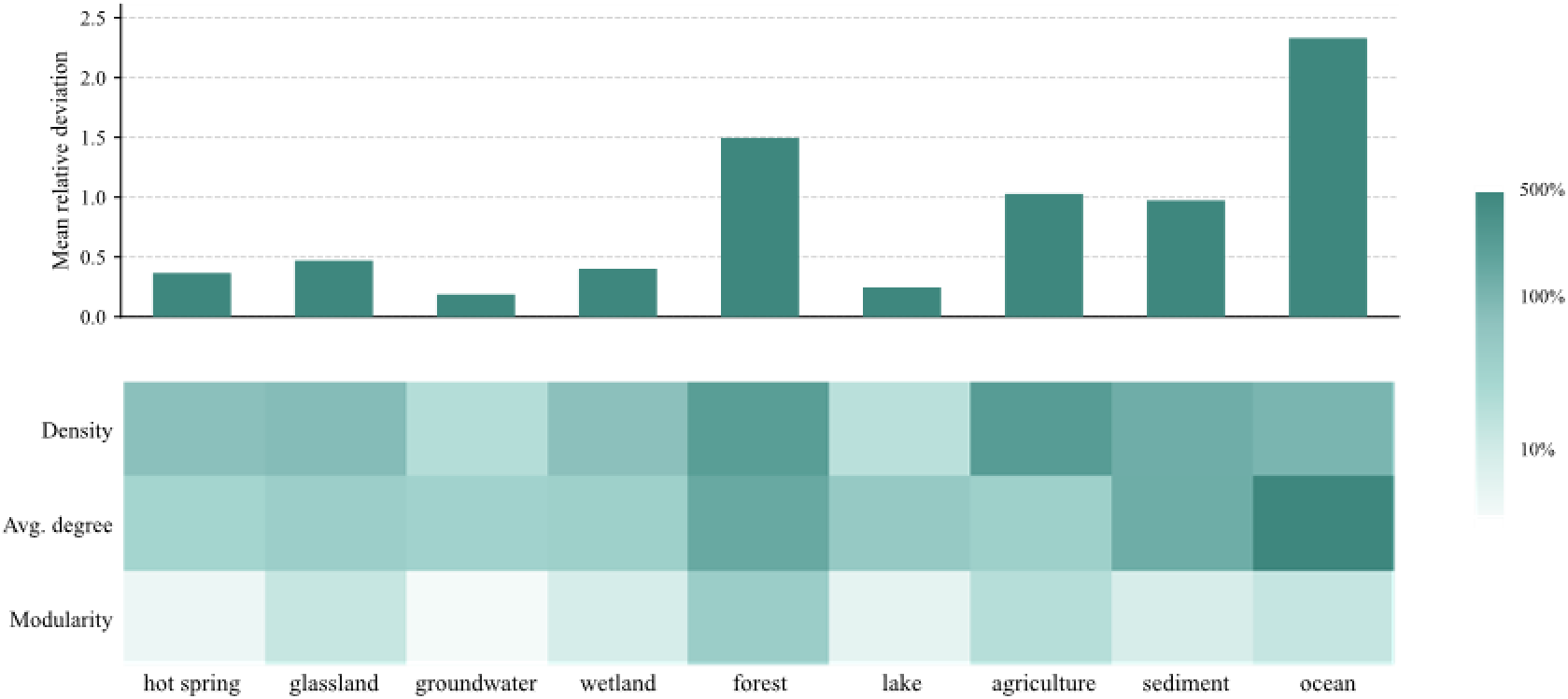
Relative deviation of generated network metrics from the real network across environments. Top, mean relative deviation (averaged over the three network metrics and six generation levels) for each environment, ordered by real sample count. Bottom, heatmap of absolute relative deviation |generated − real|/real (averaged over the six levels) for each metric and environment, displayed on a logarithmic colour scale (10–500%). Deviations are smallest in low-sample environments and largest in data-rich environments, with average degree and density accounting for most of the deviation and modularity remaining comparatively stable.

Together, Figs. 5 and 6 indicate that the fidelity of generated network topology is governed by the match between generation and training scales rather than by a fixed generation size being universally adequate. For simple, sparsely sampled communities, a few hundred generated communities already capture the co-occurrence structure. For complex communities supported by large real datasets, the same small number of generated communities systematically underfits the network, and generation sizes comparable to the available real sample count are required to faithfully reflect the real topology. In other words, the more complex the real microbial community—and the larger the real sample base supporting it—the more poorly a small, fixed amount of synthetic data approximates it, and only a matched amount of generated data can reliably reproduce the real community structure. This scale-matching requirement should be considered when interpreting network-based evaluations of generative models and when designing synthetic-data augmentation for complex habitats.

In summary, MINet outperformed MGM on the majority of the 13 metrics, including distribution similarity, biomarker retention and network topology, with significant differences on 12 of them; the only exception was environment-classification ROC-AUC, where the two models did not differ significantly. The generation-size analysis further shows that evaluation stability is metric-dependent: distribution-level metrics are robust to sample count, whereas network topology must be assessed with generation and reference sizes matched, particularly for complex, data-rich environments.

## Materials and Methods

### Construction of the microbial co-occurrence network

Microbial community data were obtained from the Earth Microbiome Project (EMP) [2]. We used the feature table emp_cr_silva_16S_123.release1.biom, processed with the standard 90-bp Deblur pipeline, together with the sample metadata file emp_qiime_mapping_release1_20170912.tsv; sequence variants were resolved using Deblur [22]. All samples had been collected, DNA-extracted, amplified, sequenced and denoised under the standardized EMP protocol, ensuring cross-environment comparability. Each feature is an exact sequence variant (ESV) taxonomically annotated against the SILVA reference database [23]. ESVs with very low total read counts were first removed, and samples were then assigned to 16 environment categories according to EMP Ontology level 3 (EMPO3; e.g., soil, water, animal gut and plant surface). A count sub-table was extracted for each category and converted to tab-separated format.

Feature filtering was performed independently within each environment to avoid spurious correlations from rare taxa. For every ESV we calculated prevalence, the fraction of samples in which it was detected, and mean relative abundance, the mean after within-sample sum normalization. An ESV was removed when its mean relative abundance was below 0.001% (1×10⁻⁵) and its prevalence below 10% simultaneously. For high-diversity environments that still retained large numbers of features, such as soils, an adaptive cap retained at most the 4,000 ESVs with the highest mean relative abundance, preserving dominant signals while limiting the quadratic growth of the correlation matrix. Feature and sample counts before and after filtering were recorded in metadata files.

Two complementary measures were computed for all ESV pairs in each filtered matrix: the Spearman rank correlation, capturing monotonic positive and negative co-occurrence, and the Bray–Curtis similarity, calculated as one minus the Bray–Curtis distance between sample-normalized abundance vectors to reflect compositional similarity [24]. Statistical significance was assessed separately for each environment with 200 permutations and 200 bootstrap iterations. Permutation tests randomly rearranged sample labels to build an empirical null distribution, and permutation P values were estimated as the proportion of permuted statistics with absolute value at least as large as the observed value, with +1 smoothing. Bootstrap resampling drew sample columns with replacement; the Welford online algorithm was used to estimate the mean and standard deviation of each edge’s statistic [25], from which a two-sided Gaussian P value was computed against zero for Spearman or against the random expectation of 0.5 for Bray–Curtis. For each measure, the larger of the permutation and bootstrap P values was taken as a conservative estimate. The two lines of evidence were combined by the Brown method [26], with the statistic T = −2(ln P_Spearman + ln P_Bray) treated as chi-square distributed with 4 degrees of freedom, and Benjamini–Hochberg (BH) correction [27] was then applied to obtain q values.

Associations with q ≤ 0.05 and |Spearman r| ≥ 0.3 were retained as significant edges, defining an undirected weighted network per environment in which each edge stored its Spearman coefficient, Bray–Curtis similarity, combined P value, q value and sign. Network deconvolution [28], with scaling parameters α = 1 and β = 0.9, was applied to the edge-weight matrix to remove indirect transitive associations, and only edges whose deconvolved weights remained above threshold were kept. The 16 environment-specific networks were then merged into a global network through a union of their significant edges; each edge received a support attribute recording the number of environments in which it was independently detected, and all edges with support ≥ 1 were retained to capture both generalist and specialist interactions. Environment-specific and global networks were exported in GraphML and edge-list formats.

All inference and statistical testing were conducted at the ESV level to preserve sequence resolution, and ESVs of the same genus were aggregated into a single node only at the interpretation and visualization stage; the weight between two genera was obtained by summing the weights of all ESV edges connecting them. For the attention prior, each inter-genus edge was assigned a composite weight W_ij = mean(r_ij) · ln(1 + n_ij), where mean(r_ij) is the mean Spearman coefficient of the underlying ESV edges and n_ij their count; the logarithmic term prevents genera containing many ESVs from being amplified solely by their edge abundance while retaining the trend that more evidence yields a higher weight. The pipeline was implemented in Python with NumPy, pandas, SciPy [29] and NetworkX; permutation and bootstrap steps were parallelized through SLURM on the high-performance computing cluster of Peking University, and all scripts and parameter settings were archived for reproducibility.

### MINet architecture and pretraining

Microbial community samples released up to June 2023 were retrieved from MGnify [30]. Following the MGM construction pipeline [10], low-quality and incomplete samples were removed, leaving 263,302 samples as the pretraining corpus. All samples were summarized at genus level, with genus names carrying the g prefix. The 9,665 distinct genera each corresponded to one token and, together with four special tokens (<PAD>, <UNK>, <SOS>, <EOS>), formed a vocabulary of 9,669.

For each sample, the relative abundance of every genus was z-score normalized using the mean and standard deviation of that genus estimated across the full corpus, and genera were ranked in descending order of z-score. After <SOS> and <EOS> were added, each sequence took the form [SOS] t₁ t₂ … tₙ [EOS], with a fixed length of 512; longer sequences were truncated and shorter ones padded with <PAD>, and the corresponding abundance values were stored in positional alignment with the tokens.

The layer-0 hidden state was the position-wise sum of three 256-dimensional embeddings followed by dropout: learned token embeddings mapping genus tokens to 256-dimensional vectors, absolute sinusoidal position encodings, and abundance embeddings produced by mapping each z-score through a two-layer MLP into 256-dimensional vectors.

The co-occurrence bias matrix was derived from the genus-level network. Edge weights were placed in a symmetric genus matrix, normalized by the maximum absolute value, and expanded to the full vocabulary according to the genus–token mapping; biases for special tokens were set to zero. The resulting matrix is a fixed prior and is not updated during training.

During the forward pass, the sequence bias was indexed by token pairs and added to the attention scores before softmax, together with causal and padding masks. The attention computation is Softmax(QKᵀ + βC), where C is the co-occurrence bias and β a sigmoid-transformed learnable scalar initialized at 0.1, shared across all eight heads; pre-layer normalization and residual connections were used at every layer. The model consists of eight Transformer blocks with hidden dimension 256, eight attention heads, feed-forward dimension 1,024, GELU activation and dropout 0.1. Token embeddings are tied with the output prediction head, and the model was implemented in PyTorch.

The pretraining objective is autoregressive next-genus prediction with cross-entropy loss that ignores padding positions (ignore_index = <PAD>). Optimization used AdamW with weight decay 0.001, excluding layer-normalization weights and biases from decay with 1,000 steps of linear warmup followed by linear decay to zero, batch size 32, 100 epochs and a gradient clipping threshold of 1.0.

Two ablated variants were trained alongside the full model: MINet (w/o bias), with the co-occurrence term removed, and MINet (w/o abundance), with the abundance MLP removed. All four models, including an MGM baseline re-implemented and retrained within our codebase rather than evaluated from its published weights, shared the same training/validation split, vocabulary, input length, optimizer, batch size and 100-epoch schedule, so that they differ only in the component under comparison. All downstream classification and generation experiments used the full checkpoint retaining the co-occurrence bias.

### Naphthalene degradation-rate prediction in synthetic consortia

The preliminary architectural validation used the synthetic-community dataset of Wang et al., which contains experimentally measured naphthalene degradation rates of 14 strains in their 2-, 3- and 4-member combinations [19]. Each consortium was treated as a sentence of microorganism tokens, using a 15-token vocabulary with tokens 1–14 for the strains and 0 for PAD. Consortia were padded to a fixed length of four, members were ordered by their initial relative abundance, and learnable position encodings were added; a binary mask marked real members against padding for attention masking and mean pooling.

Two input configurations were built: Transformer, with x = TokenEmbedding + PositionEmbedding, and Transformer (abu.), additionally including an abundance MLP that expands the scalar abundance at each position through Linear(1→64) → ReLU → Linear(64→64). In the rank-only model abundance enters solely through ordering, whereas the abundance model also receives its numerical value. Both models used two Transformer encoder layers (d_model = 64, nhead = 4, dim_ff = 128, dropout = 0.1); masked mean pooling produced a consortium-level representation passed to a regression head.

Two conventional controls were trained on 14-dimensional multi-hot membership vectors without abundance or explicit interaction modelling: Ridge regression with alpha = 1.0 as an additive linear baseline and a random forest regressor with 300 trees as a nonlinear baseline. These settings were fixed before cross-validation and shared across folds, with all other parameters kept at scikit-learn defaults; no hyperparameter selection was performed on validation folds.

Labels were z-score standardized within each fold using training-set statistics only. The Transformer models were optimized with Adam (lr = 1×10⁻³, weight_decay = 1×10⁻⁴) for 200 epochs under MSE loss. StratifiedGroupKFold (five folds, seed = 0) grouped all replicates of the same member set into one fold, so that no combination seen in training appeared in validation, while stratifying folds by quantile bins of degradation rate [31]. All four methods shared the same splits. Because R² was low and poorly discriminative, the Pearson correlation between predicted and measured rates served as the primary metric, and method differences were assessed by two-sided paired t-tests on the five fold-level correlations.

### Downstream classification tasks

Four classes of downstream tasks were used, with identical splits, preprocessing and metrics across models. (i) Ecological niche classification: MGnify samples classified at five hierarchical biome levels (Biomes 1–5) by five-fold cross-validation. (ii) Generalization to newly released samples: models trained on pre-June 2023 samples were applied without parameter updates to 75,475 MGnify samples released between June 2023 and December 2025. (iii) Cross-regional IBD diagnosis: two independent cohorts from Ireland and Canada, evaluated under three settings; RF was excluded from this task. (iv) Infant developmental-stage prediction: the longitudinal cohort of Roswall et al. [20], classified across five time points (birth, 4 months, 12 months, 3 years and 5 years).

F-max is the highest F1 score attainable over a sweep of decision thresholds, and ROC-AUC is the macro-averaged area under the receiver operating characteristic curve across classes; both range from 0 to 1, with higher values indicating better performance.

In-sample evaluation is five-fold cross-validation within the target cohort. In zero-shot transfer, a model trained without any target-region samples is applied directly to the target cohort; in fine-tuning, the zero-shot model is further adapted on a small target-region subset before evaluation.

Results are reported as mean ± SD over five folds. Model differences were tested with one-sided Welch t-tests on the five fold-level values (n = 5), with the alternative hypothesis that MINet has a higher mean; significance thresholds were P < 0.05, P < 0.01 and P < 0.001, non-significant comparisons were left unmarked, and no additional multiplicity correction was applied.

### Community generation and evaluation

Environment-conditioned generation was evaluated on 26,334 MGnify samples retrieved under nine environment labels (agriculture, forest, grassland, groundwater, hot spring, lake, ocean, sediment and wetland) [30]. Samples were summarized at genus level using the pretraining taxonomy; all 2,304 observed genera were contained in the 9,665-genus vocabulary. Training and validation partitions were shared by MINet and MGM.

Starting from the full pretrained checkpoint, the generator was fine-tuned with an environment-label token prepended to every sequence; the label token was learned during fine-tuning and constrained decoding to the target habitat. Sequences were generated by nucleus sampling with top-p = 0.9 and temperature 1.0, using identical decoding settings for both models; label tokens were excluded from the output and no genus was repeated within a community.

Because the generator emits ordered genus sequences without abundances, a dedicated reconstructor following the MGM ranking–reconstruction framework assigned abundance values [10]. It is a standalone three-layer residual network with parameters separate from the generator, taking genus presence and ranked position as input and producing softmax-normalized outputs masked to the present genera; it was trained on real (ranking, abundance) pairs using Bray–Curtis dissimilarity as the loss. No generated sequences entered its training, and it was not updated during generation, preventing abundance assignment from compensating for generator errors. Generated sequences were mapped to abundance vectors and renormalized to sum to one.

Each model generated 500 communities per environment (4,500 in total) for the primary comparison. The size-sensitivity analysis added five fixed sizes (500, 1,000, 2,000, 5,000 and 10,000) and a "same" size equal to the environment’s real training count, giving six levels.

Diversity and sparsity were computed per sample on renormalized abundance vectors: Shannon diversity (base 2) and Simpson diversity from genus proportions, and sparsity as per-sample Shannon entropy in nats using scipy.stats.entropy; all-zero samples were treated as missing. For network topology, a co-occurrence network was built independently for real and generated data, retaining genera with prevalence ≥ 1% as nodes and adding an edge only when pairwise Pearson correlation satisfied |r| ≥ 0.6 and passed BH-FDR at q < 0.05; isolated nodes were removed. Network density, average degree and modularity were computed with NetworkX; modularity used the Louvain algorithm (seed 42) [32].

Distribution similarity was measured between per-environment mean abundance vectors using cosine similarity, Spearman correlation and mean absolute error, and sequence length was summarized by mean and median. For biomarker coverage, a one-vs-rest random forest (300 trees, balanced class weights) was fit separately on real and generated data, and impurity-based feature importance selected the top 50 genera; coverage was 100 times the size of the overlap divided by 50, with identical settings for both models. Environment-classification ROC-AUC used a random forest (300 trees, balanced weights) trained on real communities and applied without modification to generated communities, with one-vs-rest ROC-AUC macro-averaged across labels.

Per-environment metric values were compared by two-sided Wilcoxon signed-rank tests across the nine paired observations, using the exact distribution at n = 9; size sensitivity was quantified by coefficients of variation across the five fixed sizes. For the ternary plots, each environment was scaled independently: an anchor range was defined around the real-network value using 1.06 times the maximum departure of all configurations, bounded by physical limits (density and modularity ≤ 1), and values were normalized to [0,1] and converted to barycentric coordinates, with arrows connecting configurations in order of increasing generation size. For the deviation analysis, absolute relative deviation |M_g − M_real|/M_real was computed per metric and level; heatmap values average the six levels, summary bars further average the three network metrics, and the heatmap colour scale is logarithmic (LogNorm).

## Acknowledgments

This work was supported by National Key R&D Program of China (2024YFA0919000) National Natural Science Foundation of China (NSFC, 32130004).

## References

1. The Human Microbiome Project Consortium. Structure, function and diversity of the healthy human microbiome. Nature, 2012, 486(7402): 207–214. DOI: 10.1038/nature11234.

[2] Thompson L R, Sanders J G, McDonald D, et al. A communal catalogue reveals Earth’s multiscale microbial diversity. Nature, 2017, 551(7681): 457–463. DOI: 10.1038/nature24621.

[3] Sunagawa S, Coelho L P, Chaffron S, et al. Structure and function of the global ocean microbiome. Science, 2015, 348(6237): 1261359. DOI: 10.1126/science.1261359.

[4] Bahram M, Hildebrand F, Forslund S K, et al. Structure and function of the global topsoil microbiome. Nature, 2018, 560(7717): 233–237. DOI: 10.1038/s41586-018-0386-6.

[5] Statnikov A, Henaff M, Narendra V, et al. A comprehensive evaluation of multicategory classification methods for microbiomic data. Microbiome, 2013, 1(1): 11. DOI: 10.1186/2049-2618-1-11.

[6] Topçuoğlu B D, Lesniak N A, Ruffin M T, et al. A framework for effective application of machine learning to microbiome-based classification problems. mBio, 2020, 11(3): e00434–20. DOI: 10.1128/mBio.00434-20.

[7] Chong H, Zha Y, Yu Q, et al. EXPERT: transfer learning-enabled context-aware microbial community classification. Briefings in Bioinformatics, 2022, 23(6): bbac396. DOI: 10.1093/bib/bbac396.

[8] Wang B, Shen Y, Fang J, et al. DeepPhylo: phylogeny-aware microbial embeddings enhanced predictive accuracy in human microbiome data analysis. Advanced Science, 2024, 11(45): 2404277. DOI: 10.1002/advs.202404277.

[9] Oh M, Zhang L. DeepMicro: deep representation learning for disease prediction based on microbiome data. Scientific Reports, 2020, 10(1): 6026. DOI: 10.1038/s41598-020-63159-5.

[10] Zhang H, Zhang Y, Kang Z, et al. MGM as a large-scale pretrained foundation model for microbiome analyses in diverse contexts. Advanced Science, 2026, 13(24): e13333. DOI: 10.1002/advs.202513333.

[11] Friedman J, Alm E J. Inferring correlation networks from genomic survey data. PLoS Computational Biology, 2012, 8(9): e1002687. DOI: 10.1371/journal.pcbi.1002687.

[12] Kurtz Z D, Müller C L, Miraldi E R, et al. Sparse and compositionally robust inference of microbial ecological networks. PLoS Computational Biology, 2015, 11(5): e1004226. DOI: 10.1371/journal.pcbi.1004226.

[13] Faust K, Raes J. Microbial interactions: from networks to models. Nature Reviews Microbiology, 2012, 10(8): 538–550. DOI: 10.1038/nrmicro2832.

[14] Rives A, Meier J, Sercu T, et al. Biological structure and function emerge from scaling unsupervised learning to 250 million protein sequences. Proceedings of the National Academy of Sciences, 2021, 118(15): e2016239118. DOI: 10.1073/pnas.2016239118.

[15] Hwang Y, Cornman A L, Kellogg E H, et al. Genomic language model predicts protein co-regulation and function. Nature Communications, 2024, 15(1): 2880. DOI: 10.1038/s41467-024-46947-9.

[16] Tackmann J, Matias Rodrigues J F, von Mering C. Rapid inference of direct interactions in large-scale ecological networks from heterogeneous microbial sequencing data. Cell Systems, 2019, 9(3): 286–296.e8. DOI: 10.1016/j.cels.2019.08.002.

[17] Aminian-Dehkordi J, Parsa M, Dickson A, et al. SIMBA-GNN: mechanistic graph learning for microbiome prediction. npj Systems Biology and Applications, 2025, 12(1): 8. DOI: 10.1038/s41540-025-00631-w.

[18] Monshizadeh M, Ye Y. Incorporating metabolic activity, taxonomy and community structure to improve microbiome-based predictive models for host phenotype prediction. Gut Microbes, 2024, 16(1): 2302076. DOI: 10.1080/19490976.2024.2302076.

[19] Wang M, Chen X, Fang Y, et al. The trade-off between individual metabolic specialization and versatility determines the metabolic efficiency of microbial communities. Cell Systems, 2024, 15(1): 63–74.e5. DOI: 10.1016/j.cels.2023.12.004.

[20] Roswall J, Olsson L M, Kovatcheva-Datchary P, et al. Developmental trajectory of the healthy human gut microbiota during the first 5 years of life. Cell Host & Microbe, 2021, 29(5): 765–776.e3. DOI: 10.1016/j.chom.2021.02.021.

[21] Bäckhed F, Roswall J, Peng Y, et al. Dynamics and stabilization of the human gut microbiome during the first year of life. Cell Host & Microbe, 2015, 17(5): 690–703. DOI: 10.1016/j.chom.2015.04.004.

[22] Amir A, McDonald D, Navas-Molina J A, et al. Deblur rapidly resolves single-nucleotide community sequence patterns. mSystems, 2017, 2(2): e00191–16. DOI: 10.1128/mSystems.00191-16.

[23] Quast C, Pruesse E, Yilmaz P, et al. The SILVA ribosomal RNA gene database project: improved data processing and web-based tools. Nucleic Acids Research, 2013, 41(D1): D590– D596. DOI: 10.1093/nar/gks1219.

[24] Bray J R, Curtis J T. An ordination of the upland forest communities of southern Wisconsin. Ecological Monographs, 1957, 27(4): 325–349. DOI: 10.2307/1942268.

[25] Welford B P. Note on a method for calculating corrected sums of squares and products. Technometrics, 1962, 4(3): 419–420. DOI: 10.1080/00401706.1962.10490022.

[26] Brown M B. A method for combining non-independent, one-sided tests of significance. Biometrics, 1975, 31(4): 987–992. DOI: 10.2307/2529826.

[27] Benjamini Y, Hochberg Y. Controlling the false discovery rate: a practical and powerful approach to multiple testing. Journal of the Royal Statistical Society: Series B, 1995, 57(1): 289–300. DOI: 10.1111/j.2517-6161.1995.tb02031.x.

[28] Feizi S, Marbach D, Médard M, et al. Network deconvolution as a general method to distinguish direct dependencies in networks. Nature Biotechnology, 2013, 31(8): 726–733. DOI: 10.1038/nbt.2635.

[29] Virtanen P, Gommers R, Oliphant T E, et al. SciPy 1.0: fundamental algorithms for scientific computing in Python. Nature Methods, 2020, 17(3): 261–272. DOI: 10.1038/s41592-019-0686-2.

[30] Mitchell A L, Almeida A, Beracochea M, et al. MGnify: the microbiome analysis resource in 2020. Nucleic Acids Research, 2020, 48(D1): D570–D578. DOI: 10.1093/nar/gkz1035.

[31] Pedregosa F, Varoquaux G, Gramfort A, et al. Scikit-learn: machine learning in Python. Journal of Machine Learning Research, 2011, 12: 2825–2830.

[32] Blondel V D, Guillaume J L, Lambiotte R, et al. Fast unfolding of communities in large networks. Journal of Statistical Mechanics: Theory and Experiment, 2008, 2008(10): P10008. DOI: 10.1088/1742-5468/2008/10/P10008.

