## Supplemental Data for "MINet: a microbiome foundation model integrating microbial co-occurrence priors with abundance information"

Supplementary Table 1 | Mean ± SD of the 13 evaluation metrics across nine environments.

| Category | Metric | Real | MGM | MINet | Sig. |
|---|---|---|---|---|---|
| Diversity Fidelity | Shannon | 3.77 ± 0.78 | 0.66 ± 0.25 | 6.55 ± 0.85 | ** |
| | Simpson | 0.79 ± 0.09 | 0.23 ± 0.09 | 0.94 ± 0.07 | ** |
| | Sparsity | 2.62 ± 0.54 | 0.46 ± 0.17 | 4.54 ± 0.59 | ** |
| Network Structure | Density | 0.0091 ± 0.0036 | 0.0469 ± 0.0196 | 0.0131 ± 0.0059 | ** |
| | Average Degree | 4.33 ± 2.11 | 28.83 ± 12.83 | 11.61 ± 8.10 | * |
| | Modularity | 0.87 ± 0.07 | 0.69 ± 0.16 | 0.81 ± 0.09 | ** |
| Distribution similarity | Cosine | – | 0.16 ± 0.09 | 0.57 ± 0.21 | ** |
| | Spearman ρ | – | 0.63 ± 0.06 | 0.71 ± 0.07 | * |
| | MAE($10^{-4}$) | – | 1.83 ± 0.085 | 1.07 ± 0.25 | ** |
| Classification and ecological validity | ROC-AUC | – | 0.918 ± 0.047 | 0.892 ± 0.077 | ns |
| | Biomarker Coverage (%) | – | 27.3 ± 10.4 | 42.0 ± 11.6 | * |
| Sequence length | Mean Length | 140.3 ± 39.4 | 92.0 ± 35.7 | 148.7 ± 10.9 | ** |
| | Median Length | 122.0 ± 47.8 | 88.2 ± 43.3 | 156.2 ± 2.7 | ** |

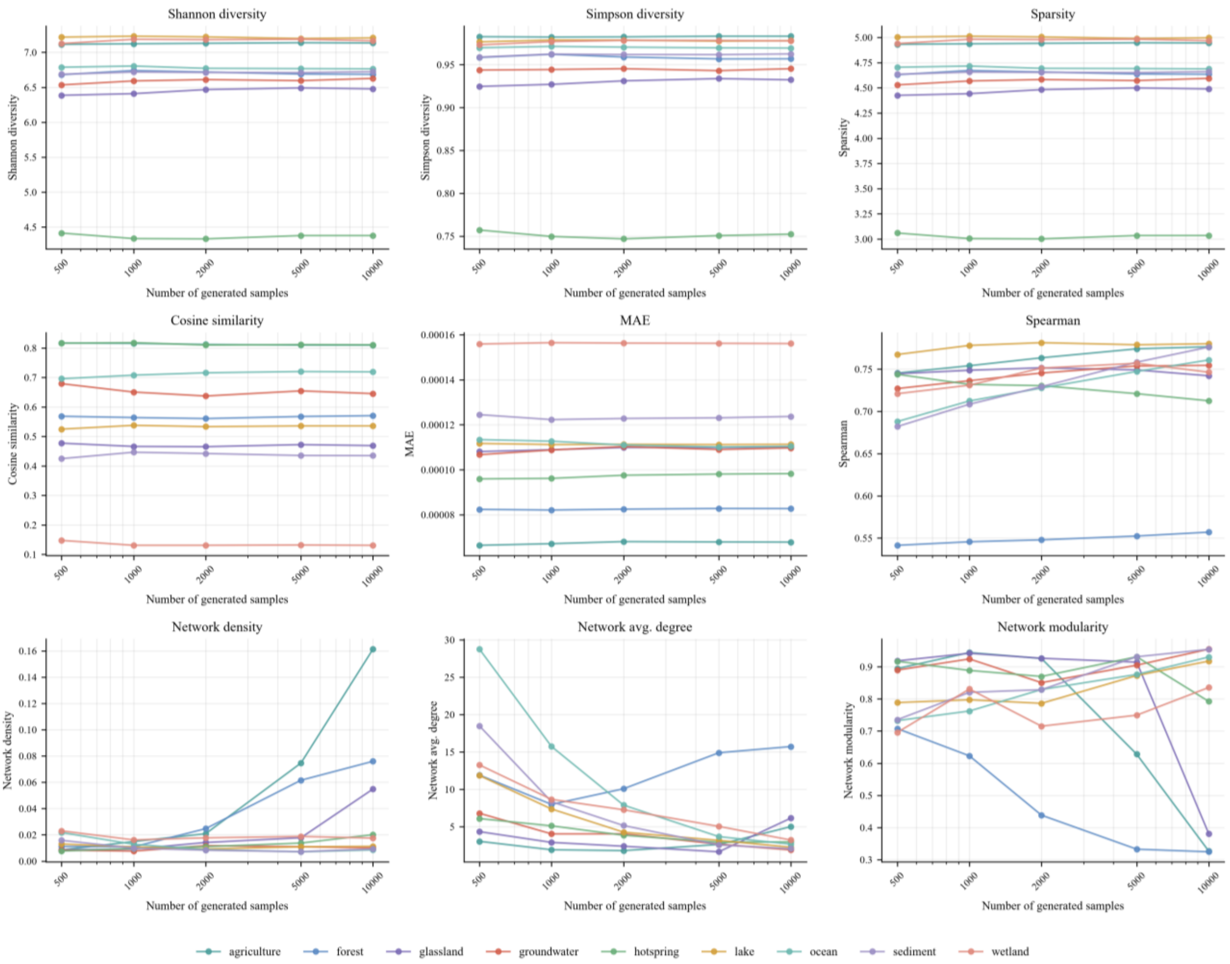


Supplementary Fig. 1 | Variation of MINet evaluation metrics with the number of generated samples. Each subplot shows the value of a metric at five fixed generation sizes, 500, 1,000, 2,000, 5,000 and 10,000 (horizontal axis on a logarithmic scale), and the nine coloured curves correspond to the nine environment labels. The first two rows show distribution-level metrics (Shannon diversity, Simpson diversity, sparsity, cosine similarity, MAE, Spearman), for which the per-label curves are nearly horizontal (mean coefficients of variation all < 2%). The last row shows co-occurrence network metrics (density, average degree, modularity), whose apparent fluctuation occurs mainly where generation and training scales are severely mismatched, whereas the metrics approach the real data when the two scales are comparable. This fluctuation reflects the inherent sample-size dependence of the correlation-network construction (edges filtered by Pearson correlation with BH-FDR significance), rather than a change in generated-data quality.
